# A high-throughput assay for the measurement of Ca^2+^-oscillations and insulin release from uniformly sized β-cell spheroids

**DOI:** 10.1101/2025.05.25.655695

**Authors:** Stijn Robben, Patricia Davidson, Rita S. Rodrigues Ribeiro, Thomas Voets

## Abstract

Diabetes mellitus is a rapidly growing global health challenge, necessitating the development of more effective anti-diabetic therapies, including drugs that improve insulin release from pancreatic β-cells. Traditional high-throughput screening methods typically rely on 2D β-cell cultures, but such cultures do not mimic the 3D organisation and cell-to-cell communication of β-cells in pancreatic islets of Langerhans. Existing 3D β-cell culture models are hindered by high costs, technical complexity, and limited compatibility with high-throughput screening platforms. In this work, we developed an approach for generating 19 homogeneously shaped pancreatic β-cell spheroids in each well of a 96-well plate, using micropatterned polyethylene glycol (PEG)-based hydrogels and MIN6 insulinoma cells. The uniform shape and positioning of the individual spheroids enabled the simultaneous, real-time imaging of Ca^2+^ signals in up to 1824 independent spheroids in response to glucose and various test compounds. Using this approach, we show that increasing glucose causes concentration-dependent Ca^2+^ oscillations in individual spheroids, that these Ca^2+^ oscillations are sensitive to modulators of ATP-sensitive K^+^ channels, and that the frequency of Ca^2+^ oscillations correlates with insulin secretion. Finally, we demonstrate that the neurosteroid pregnenolone sulphate, an agonist of the cation channel TRPM3, increases the frequency of glucose-induced Ca^2+^ oscillations and enhances insulin release; the TRPM3 antagonist isosakuranetin inhibited these responses. In conclusion, we established a cost-effective and scalable 3D β-cell platform for high-throughput screening of insulin release-modifying compounds, with potential applications in drug development and personalized medicine for the management of diabetes mellitus.

## Introduction

As one of the fastest growing diseases, diabetes mellitus is estimated to affect 783.2 million adults (20-79 years) worldwide by 2045^1^. Characterised by insulin resistance and/or abnormal insulin secretion, this disease leads to increased mortality, blindness, kidney failure and an overall decreased quality of life^2^. In 2019, it was estimated that 4.2 million adults aged 20-79 years died from diabetes-related complications, with almost half of these deaths (1.9 million) estimated to have occurred in adults below the age of 60^3^. These premature deaths combined with disability due to diabetes and absenteeism from work contribute to an astronomical economic impact, estimated to a total of 1.31 trillion USD in 2019 alone^4^. Hence, the development of effective anti-diabetic drugs, including improved insulin secretagogues, has been a priority for the pharmaceutical sector.

*In vitro* cell-based high-throughput screening methods often form a starting point for the development of new drugs^5, 6^. For the discovery of insulin secretagogues, currently used screening methods include cytosolic calcium measurements and insulin secretion assays, primarily performed on pancreatic β-cells cultured in a 2D monolayer^7^. However, 2D cell models can be suboptimal predictors of medical outcomes in clinical trials due to the lack of physiological or morphological relevance to *in vivo* conditions^8, 9,10^. This is particularly relevant for pancreatic β-cells, which are clustered *in vivo* in the islets of Langerhans. In these highly metabolically active spherical clusters of endocrine cells, dispersed throughout the exocrine tissue of the pancreas, β-cells exhibit coordinated activity in response to glucose, which leads to islet calcium waves and pulsatile insulin release. To date, only few studies have explored the application of 3D β-cell culture for developing high-throughput-compatible screening methods^11, 12^. The main challenges remain the high costs, technical complexity, time consumption, and issues with reproducibility due to variability and lack of standardization^13^. Furthermore, the limited compatibility of these models with high-throughput read-out methods poses a significant barrier to their integration into industrial applications, complicating the development of a standardized and reproducible assay that meets all necessary requirements.

In this work, we introduce a novel approach which allows the precisely patterned creation of uniform 3D pancreatic β-cell spheroids within a 96-well plate, utilizing micropatterned polyethylene (PEG)-based hydrogels. Due to the unique design and properties of the plates, we manage to achieve simultaneous real-time monitoring of insulin release-associated Ca^2+^-fluorescence signals at the individual spheroid level, yielding a total of up to 1824 unique measurements per plate (19 spheroids per well), and directly correlate these signals with insulin release. This method solely relies on conventional cell culture techniques, with results obtainable within 24 hours after cell seeding. As a proof of principle, we illustrate the concentration-dependent effects of modulators of ATP-sensitive K^+^ (K-ATP) and TRPM3 channels on glucose-induced Ca^2+^ oscillations and insulin release.

## Results

### Hydrogel design

To facilitate spheroid formation while ensuring compatibility with high-throughput calcium screening, 316L stainless-steel molds were designed to produce gels conforming to the dimensions of the well of a 96-well plate and contain 19 identical microwell structures into which the cells will sediment after seeding (Fig. 1A-C). The molds were used to stamp multiple gels in a single polymerization step, resulting in a polyethylene glycol (PEG) based hydrogel covered glass-bottom 96 well plate (Fig. 1D). The uniformity in the shape and dimensions of the fabricated microstructured hydrogels was assessed by quantifying the contact point diameters. These contact points (Fig. 1E - top) correspond to regions where the molds make direct contact with the glass substrate during polymerization, inhibiting hydrogel formation in those areas. The mean contact point diameter was determined to be 183 µm ± 23 µm (mean ± SD), with no statistically significant variation observed between samples (p > 0.05) (Fig. 1E - bottom).

**Figure 1.**
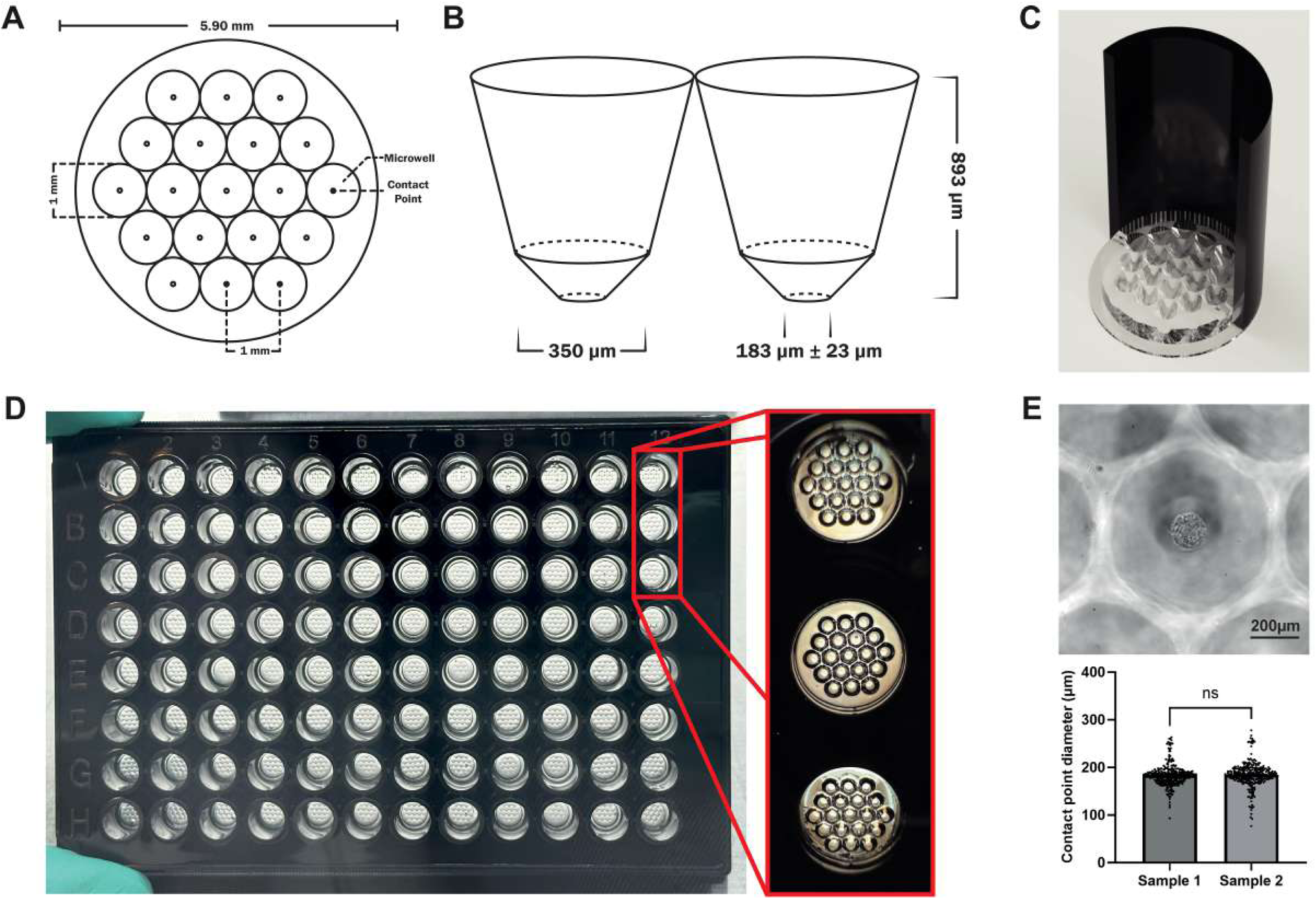
Hydrogel design. **(A)** Schematic top-view representation of the hydrogel, illustrating a honeycomb arrangement of 19 microwells per gel. **(B)** Schematic cross-sectional depiction of the microwells (not to scale). **(C)** CAD rendering of the hydrogel positioned within the well of a glass-bottom 96-well plate. **(D)** Entire glass-bottom 96-well plate with microstructured PEG-based hydrogel in each well. **(E)** Microwell with Mouse Embryonic Fibroblasts attached to the contact point, which is a result of the molds being in contact with the glass bottom during polymerization (top). Mean contact point diameter for two independently moulded samples. N = 16 wells (304 contact points) per sample. Statistical difference was analyzed by one-way ANOVA and Tukey’s post-hoc test where ns = not significant.

### Protocol

After determining that the microwells of our hydrogels within the 96-well plate are of consistent shape, we focussed on developing a protocol to assess intracellular calcium signalling in the 3D pancreatic β-cell spheroids. Figure 2A illustrates the established timeline including the tissue formation, loading of the tissues with a Ca^2+^-sensitive dye and data acquisition. Within 16 hours post-seeding of the MIN6 β-cells onto the hydrogels, 3D tissues had formed within the microwells (Fig. 2B) and changes in the Ca^2+^-influx associated fluorescent signal of individual spheroids could be measured in real-time using a fluorescence plate imager (Figure 2C). The contact points exposing the glass substrate facilitate tissue adhesion, preventing spheroid loss during washing steps and liquid deposition or aspiration by the µCELL FDSS. Following all experimental manipulations, 75.0% of the wells retained all 19 spheroids, with no wells losing more than three spheroids despite the extensive washing steps.

**Figure 2.**
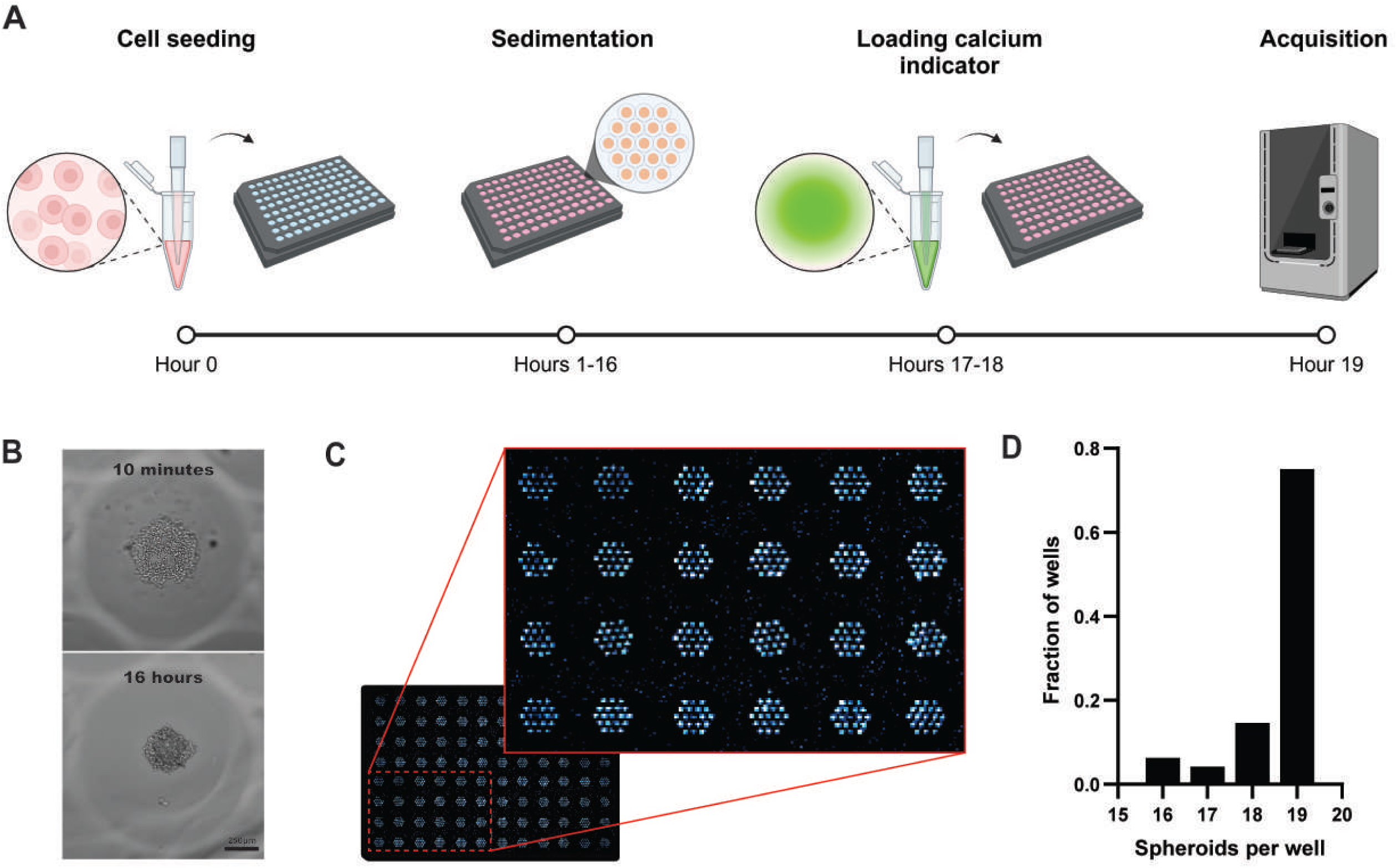
Cell seeding and data acquisition. **(A)** Schematic representation of protocol timeline. 200 μL of the MIN6 pancreatic β-cell suspension is added to each well of the 96-well plate at hour 0. Over the next 16 hours, the cells undergo sedimentation into the microwells and spontaneously self-assemble into spheroids. Two hours prior to data acquisition, 10 μM of Cal-520 is introduced for 1 hour, followed by washing and incubation in a low-glucose KREBS solution for 30 minutes (hours 17–18). At hour 19, the plates are transferred to the μCELL FDSS for fluorescence-based data acquisition. **(B)** Representative images of MIN6 β-cells following sedimentation within a microwell, captured 10 minutes post-seeding (top) and 16 hours post-seeding (bottom). **(C)** Representative fluorescence image of a section of a 96-well plate containing the Cal-520 AM-loaded MIN6 β-cell spheroids acquired using the µCELL FDSS fluorescence plate imager, distinctly showing individual spheroids arranged in a honeycomb pattern within the microwell array. **(D)** Histogram showing the number of spheroids still present per well at the time of the control stimulus. N = 48 wells on two independent occasions.

### Increased levels of extracellular glucose induce Ca^2+^-oscillatory behaviour

Next, we evaluated the spheroid Ca^2+^ signal response to increases in extracellular glucose concentration (Figure 3). At the basal glucose concentration (3 mM), the signal remained stable, free of any oscillatory behaviour. However, periodic oscillations emerged at extracellular glucose concentrations exceeding 5 mM, with increased frequency at higher glucose concentrations (Fig 3A,B), whereas no oscillations were observed in 2D cultures of the same cells (Fig.3 - Figure supplement 1). To quantify the glucose-induced Ca²⁺ responses, we determined two main oscillatory parameters: the frequency and the area under the peaks (AuP), as illustrated in Figure 3A. Both the frequency and AuP increased in a concentration-dependent manner with higher glucose, with concentrations for half-maximal activation (EC□□ values) of 7.4 ± 0.4 mM and 7.9 ± 0.4 mM for frequency and AuP, respectively. Figure 3C displays the fluorescence traces of all individual spheroids within a singe well, prior to baseline subtraction, and the average of those signals. Figure 3D presents a comparison of the average Ca²⁺ fluorescence signal recorded from a well containing 3D-cultured MIN6 β-cell spheroids and a well containing MIN6 cells cultured under conventional 2D conditions.

**Figure 3.**
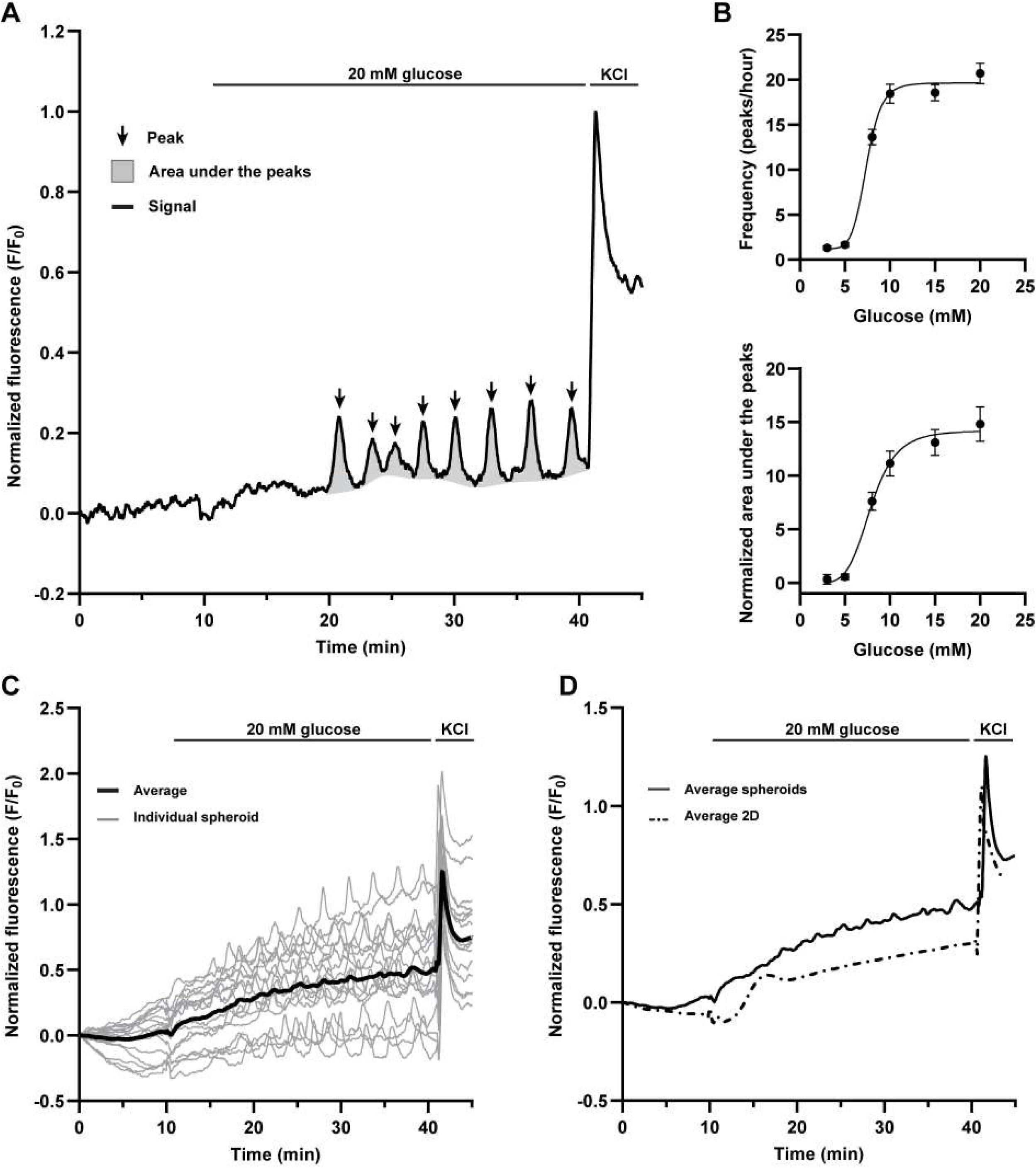
Oscillatory behaviour of Ca^2+^ signal is modulated by extracellular glucose concentrations. **(A)** Representative Ca²⁺ response of a single 3D MIN6 β-cell spheroid to an increase in extracellular glucose concentration (20 mM), highlighting the visual representations of the peaks and the area under the peaks. The spheroids were initially exposed to basal extracellular glucose levels (3 mM) for 10 minutes, followed by an increase in extracellular glucose (20 mM) for 30 minutes. The protocol ended with a control stimulus of 50 mM KCl. **(B)** Mean frequency and the normalized area under the peaks of the fluorescent signal during the 30 minutes of exposure to increased concentrations of extracellular glucose. Peak threshold was set to 3 times the signal’s standard deviation during the initial 10 minutes of exposure to basal levels of extracellular glucose. Error bars represent the standard error of the mean (SEM). N > 50 spheroids per concentration. Signal values for each spheroid were normalized to the amplitude of the control stimulus. **(C)** Overview of Ca²⁺ responses of each individual MIN6 β-cell spheroid within a single well to an increase in extracellular glucose concentration (20 mM). To preserve the absolute response dynamics and inter-spheroid variability, no baseline correction was applied to the displayed traces. Gray lines represent individual spheroids, while the black line denotes the mean response across all spheroids. **(D)** Comparison of the mean Ca²⁺ fluorescence response of all individual MIN6 β-cell spheroids within a single well (solid line) and 2D-cultured MIN6 cells (dashed line) following stimulation with 20 mM extracellular glucose.

### K_ATP_-agonist diazoxide inhibits Ca^2+^-oscillatory response to extracellular glucose

It is well-established that glucose-induced insulin release *in vivo* depends on the inhibition of K-ATP channels, which close when cytosolic ATP levels rise in response to glucose in flux and metabolism. To assess the dependence of glucose-induced calcium oscillations in the MIN6 β-cell spheroids on K-ATP channel closure, we exposed spheroids to different concentrations of diazoxide, a K-ATP channel opener, prior to stimulation with 10 mM glucose. Diazoxide caused a concentration-dependent inhibition of the glucose-induced oscillations (Fig. 4A). Both the frequency and AuP decreased with increasing diazoxide, with concentrations for half-maximal inhibition (IC_50_ values) of 2.9 ± 0.1 μM and 1.9 ± 1.71 μM, respectively.

**Figure 4.**
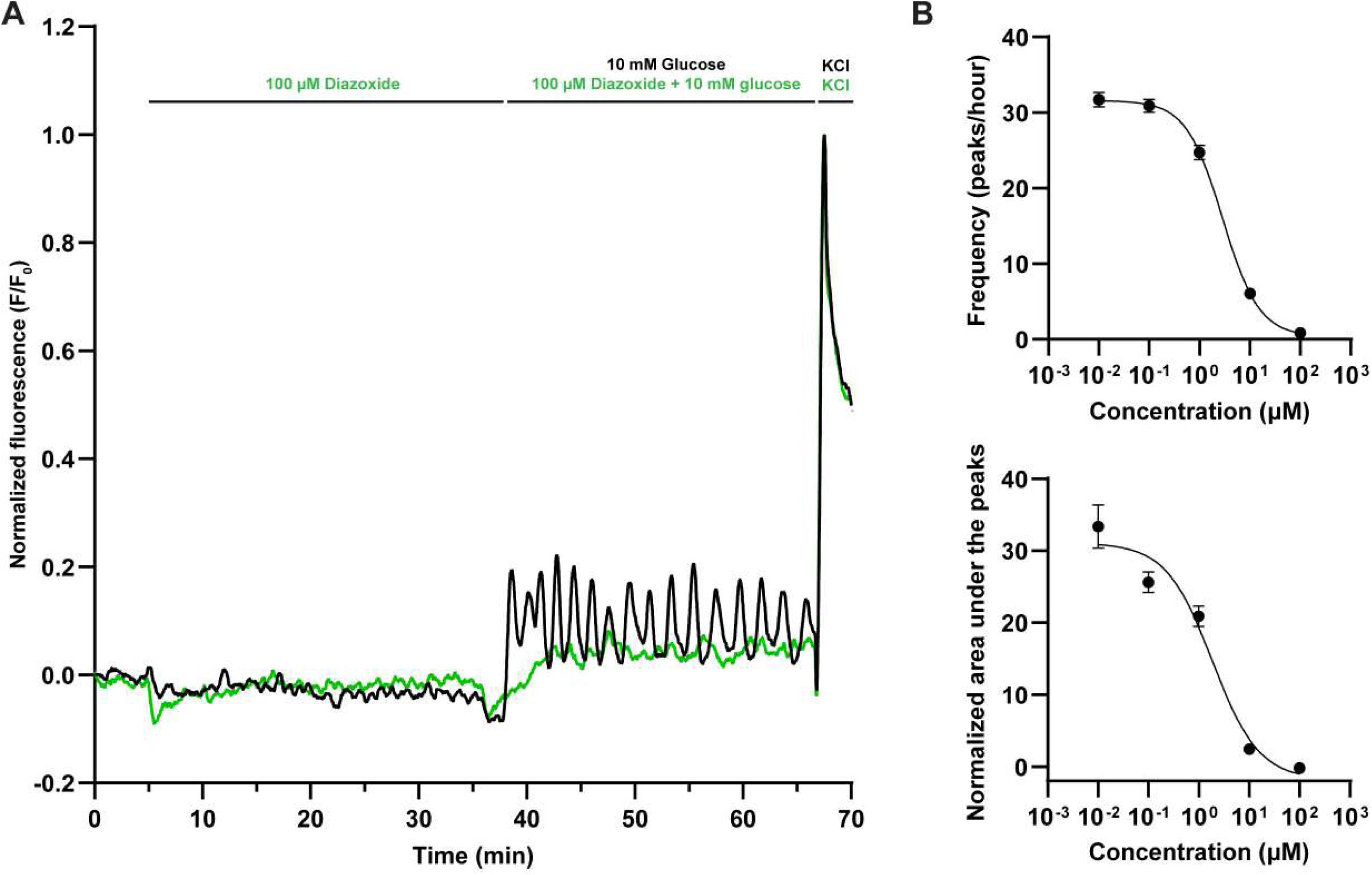
K_ATP_-agonist diazoxide inhibits oscillatory response to extracellular glucose. **(A)** Characteristic Ca^2+^ responses to 10 mM extracellular glucose concentrations of individual MIN6 β-cell spheroids with and without 30 minutes of prior exposure to 100 μM diazoxide. The spheroids were initially exposed to basal extracellular glucose levels for 5 minutes after which the spheroids were exposed to varying concentrations of diazoxide for 30 minutes, followed by a 30 minute period of extracellular glucose concentrations of 10 mM until a control stimulus of 50 mM KCl. Each signal was normalized over the amplitude of the control stimulus. **(B)** Average of the frequency and the normalized area under the peaks of the fluorescent signal during the 30 minutes of exposure to increased concentrations of extracellular glucose. Peak threshold was set to 3 times the signal’s standard deviation during the initial 10 minutes of exposure to basal levels of extracellular glucose. Error bars represent the standard error of the mean (SEM). N > 40 spheroids per concentration.

### K_ATP_-antagonists induce Ca^2+^-oscillatory behaviour

K_ATP_-channel antagonists, such as the sulfonylureas and meglitinides, are widely used to increase insulin release in patients with type 2 diabetes. To test the efficacy of K_ATP_-channel antagonists in our MIN6 β-cell spheroid assay, we exposed spheroids in 3 mM glucose to increasing levels of glimepiride for 30 minutes, and then increased extracellular glucose to 10 mM. As illustrated in Figure 5A, at a concentration of 100 nM, the potent sulfonylurea antagonist glimepiride induced robust calcium oscillations. Glimepiride increased both the frequency and AuP in a concentration-dependent manner, with EC□□ values of 3.3 ± 0.6 nM and 3.4 ± 1.5 nM, respectively. Similar results, albeit with a different concentration dependence, were obtained with the less potent, first-generation sulfonylurea antagonist tolbutamide (EC□□ values of 6.4 ± 6.1 μM for the frequency and 4.3 ± 7.7 μM for the AuP) and with the meglitinide antagonist nateglinide (EC□□ values of 44.5 ± 1.23 nM for the frequency and 67.4 ± 0.623 nM for the AuP). Taken together, these results illustrate the ability of the MIN6 β-cell spheroid assay to quantitatively measure the effects of K_ATP_ modulators on intracellular Ca^2+^ dynamics.

**Figure 5.**
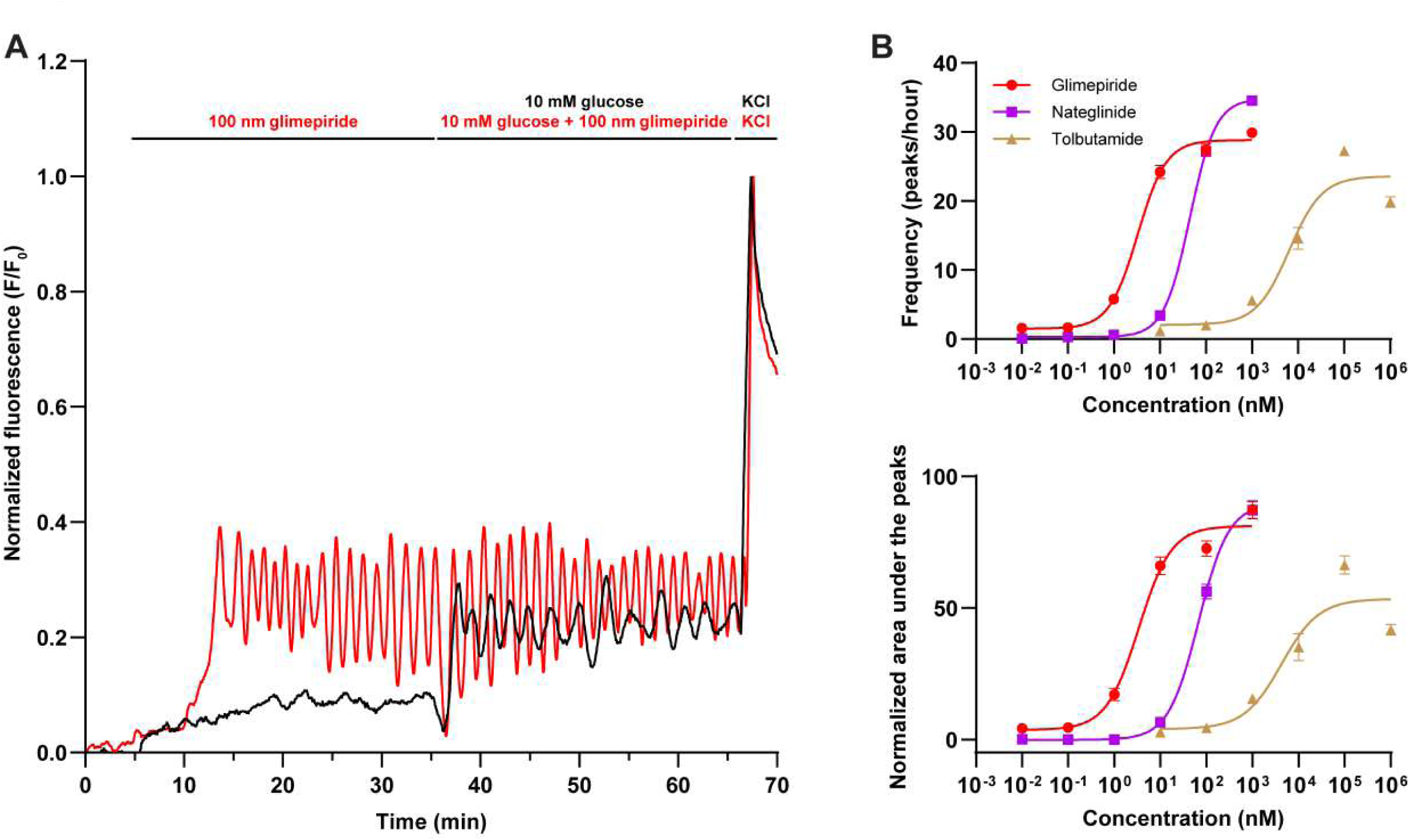
K_ATP_-antagonists induce Ca^2+^ oscillatory behaviour with distinct potencies. **(A)** Characteristic Ca^2+^ responses of individual MIN6 β-cell spheroids to 10mM of extracellular glucose with and without prior exposure to 100 nM glimepiride. The spheroids were initially exposed to basal extracellular glucose levels for 5 minutes after which the spheroids were exposed to varying levels of glimepiride for 30 minutes, followed by a 30 minute period of glimepiride + extracellular glucose concentrations of 10mM until a control stimulus of 50 mM KCl. Each signal was normalized over the amplitude of the control stimulus. **(B)** Average of the frequency and the normalized area under the peaks of the fluorescent signal during the 30 minutes of exposure to various concentrations of glimepiride, nateglinide and tolbutamide. Peak threshold was set to 3 times the signal’s standard deviation during the initial 10 minutes of exposure to basal levels of extracellular glucose. Error bars represent the standard error of the mean (SEM). N > 50 spheroids per concentration.

### K_ATP_-channel modulators impact insulin secretion by MIN6 spheroids

To determine whether Ca^2+^ oscillation in the MIN6 β-cell spheroids translate into insulin secretion, we compared calcium signals and insulin release in three conditions: 8 mM extracellular glucose in the presence of either vehicle, 100 nM glimepiride or 100 µM diazoxide. After measuring calcium signals during 60 minutes (Fig. 6A,B), supernatant was aspirated from the wells and the insulin content of aspirated samples was quantified using ELISA (Fig. 6C). The results indicate that treatment with 8 mM glucose caused calcium oscillations and increased insulin secretion. Inclusion of glimepiride (100 nM) not only increased the frequency of oscillations, but also robustly enhanced insulin release. Oppositely, diazoxide (100 μM) abolished calcium oscillations, and reduced the insulin in the supernatant to the level of non-stimulated spheroids. These data demonstrate that the frequency of evoked calcium oscillations in MIN6 β-cell spheroids positively correlates with insulin release.

**Figure 6.**
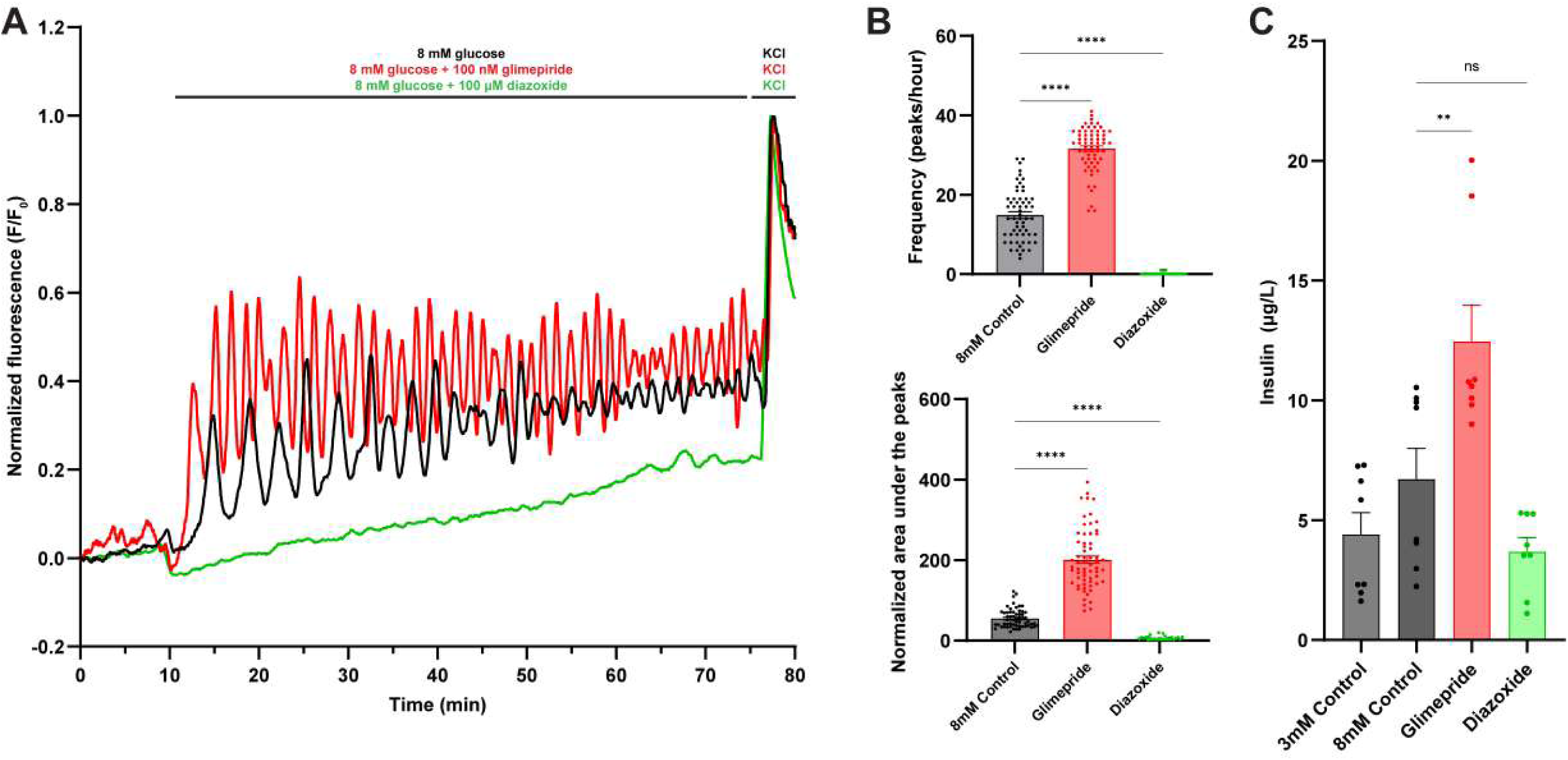
Glimepiride and diazoxide impact glucose dependent insulin release by MIN6 spheroids. **(A)** Characteristic Ca^2+^ responses of individual MIN6 β-cell spheroids. The spheroids were initially exposed to basal extracellular glucose levels for 10 minutes after which the spheroids were exposed to either 8 mM of extracellular glucose, or 8 mM of extracellular glucose supplemented with 10 nM glimepiride or 10 µM diazoxide followed by a control stimulus of 50 mM KCl. Each signal is normalized over the maximum value of the control stimulus **(B)** Average of the frequency and the normalized area under the peaks of the fluorescent signal during the 60 minutes of exposure to the various modulators. Peak threshold was set to 3 times the signal’s standard deviation during the initial 10 minutes of exposure to basal levels of extracellular glucose. Error bars represent the standard error of the mean (SEM). N > 50 spheroids per concentration. **(C)** Corresponding insulin concentrations present in the supernatant of the wells at the 1-hour timepoint quantified using ELISA. Data is presented as means ± SEM (n = 4 wells per condition). Statistical difference was analyzed by one-way ANOVA and Tukey’s post-hoc test where ns = not significant, ** p < 0.01, **** p < 0.0001.

### Pregnenolone sulfate (PS) induces Ca^2+^-oscillatory behaviour and increases insulin secretion

To evaluate whether other signalling pathways can influence Ca²⁺ signals in MIN6 β-cell spheroids, we tested the effect of pregnenolone sulphate (PS), an endogenous neurosteroid and agonist of the cation channel TRPM3. At a concentration of 50 μM, PS strongly potentiated glucose-induced calcium oscillatory behaviour (Fig. 7A,B), and this was associated with enhanced insulin release (Fig. 7C). To evaluate whether the PS-induced increase in calcium oscillations and insulin release is dependent on TRPM3, we performed additional experiments in which PS was applied in the absence or presence of the potent TRPM3 antagonist isosakuranetin (ISN; 10 μM). These experiments revealed that ISN partly reversed effects of PS on calcium oscillations and insulin release, indicating that the positive effects of PS on insulin release are at least partly mediated by TRPM3. To assess the robustness and assay quality of the calcium imaging measurements in MIN6 β-cell spheroids, the Z-factor was calculated and determined to be 0.6.

**Figure 7.**
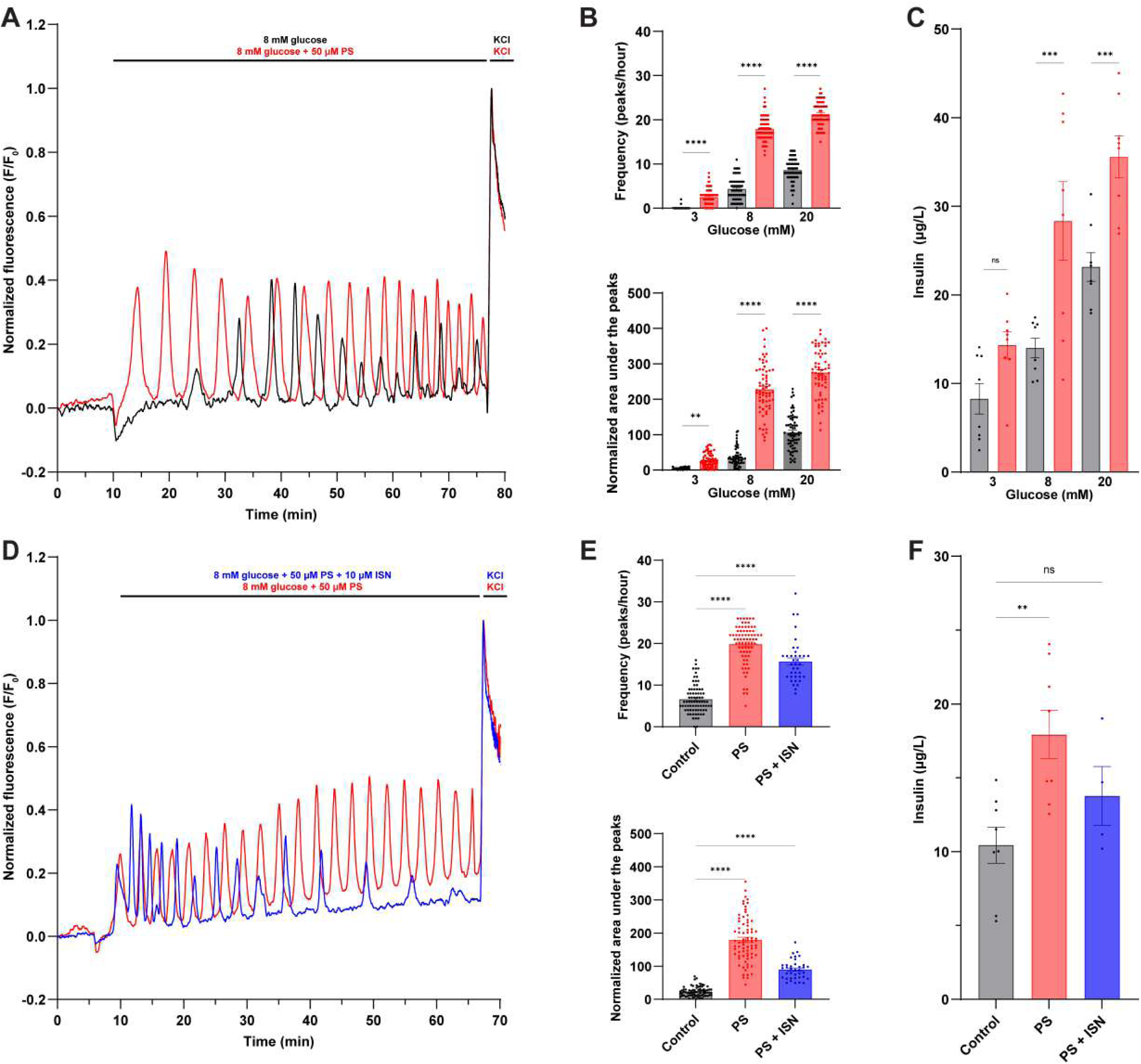
PS induces Ca^2+^-oscillatory behaviour and increases insulin secretion. **(A)** Characteristic Ca²⁺ responses of individual MIN6 β-cell spheroids exposed to an extracellular glucose concentration of 8 mM, with and without 50 μM PS. Spheroids were initially exposed to a baseline extracellular glucose concentration of 3 mM for 10 minutes, followed by a 1-hour incubation with varying glucose concentrations, with or without 50 μM PS. After 60 minutes, 15 μL of supernatant was collected for insulin level analysis, concluding with a control stimulus of 50 mM KCl. **(B)** Quantitative analysis of Ca²⁺ oscillation parameters. The frequency and normalized area under the peaks were analysed for spheroids exposed to glucose concentrations of 3 mM, 8 mM, and 20 mM, with or without 50 μM PS. Peak threshold was set at 3 times the signal’s standard deviation during the initial 10 minutes of exposure to basal levels of extracellular glucose. **(C)** Corresponding insulin concentrations present in the supernatant of the wells at the 1-hour timepoint quantified using ELISA. Data is presented as means ± SEM (n = 4 wells per condition). **(D)** Characteristic smoothed Ca²⁺ responses of individual MIN6 β-cell spheroids exposed to an extracellular glucose concentration of 8 mM + 50 μM PS, with and without 10 μM isosakuranetin (ISN). Spheroids were initially exposed to a baseline extracellular glucose concentration of 3 mM for 5 minutes, followed by a 1-hour incubation with 8 mM of extracellular glucose + 50 μM PS with and without 10 μM ISN. After 60 minutes, 15 μL of supernatant was collected for insulin level analysis, concluding with a control stimulus of 50 mM KCl. **(E)** Quantitative analysis of Ca²⁺ oscillation parameters. The frequency and normalized area under the peaks were analysed for spheroids exposed to a glucose concentration of 8 mM with or without 50 μM PS, supplemented with or without 10 μM ISN. Peak threshold was set at 5 times the signal’s standard deviation during the initial 10 minutes of exposure to basal levels of extracellular glucose. **(F)** Corresponding insulin concentrations present in the supernatant of the wells at the 1-hour timepoint quantified using ELISA. Data is presented as means ± SEM (n = 4 wells per condition). Statistical difference was analyzed by one-way ANOVA (E&F) or two-way ANOVA (B&C) and Tukey’s post-hoc test where ns = not significant, ** p < 0.01, *** p < 0.001, **** p < 0.0001.

## Discussion

In this study, we describe a platform to create homogeneously sized β-cell spheroids in a 96-well plate, and illustrate how these can be used to screen for small-molecule modulators of glucose-induced Ca^2+^ oscillations and insulin release. In particular, within 24 hours after cell seeding, 1824 spheroids (19 per well) uniformly sized spheroids are formed, and their Ca^2+^ signals can be measured simultaneously using commercial fluorescent plate imaging systems. We found that β-cells within each spheroid exhibit synchronized Ca^2+^ kinetics, resulting in spheroid calcium oscillations, whereas the 19 spheroids within the same well show similar but temporally-independent Ca^2+^ signals. Analysis of insulin levels in individual wells demonstrated the correlation between oscillatory frequency and insulin secretion.

Pancreatic β-cells in 3D cell culture have been used in earlier studies, with numerous reports highlighting the advantages of spheroid-based approaches to emulate *in vivo* pancreatic islets^12, 14, 15^. Several methods have been described for the generation of these spheroids, including the use of the hanging-drop method, microwell systems, scaffold-based culture and ultra-low adherence plates^16^. From these, microwell systems have emerged as the most cost-effective and scalable option for producing uniform and functional β-cell spheroids^17^. One earlier study reported the use of microwell method combined with microfluidics to form β-cell spheroids, and applied Ca^2+^ imaging to a single spheroid, observing synchronized Ca^2+^ oscillations throughout the spheroid^12^. However, large-scale, parallel Ca^2+^ imaging of 3D β-cell tissues, with comparable throughput as in 2D screening methods, has not been achieved. We present a method that addresses this gap by providing a platform enabling high-throughput screening performance, as reflected by a Z′-factor of 0.6, indicating excellent assay quality for drug screening, together with detailed analysis of Ca2+ influx oscillations in pancreatic β-cells cultured within a highly reproducible 3D environment. This approach enables screening for insulin secretagogues in a 3D environment, enhancing the physiological relevance and increasing the predictive power of drug discovery efforts.

The β-cell spheroids develop by self-organization of MIN6 cells guided by the specific design of our micropatterned hydrogel, which we here implemented within the individual wells of a 96-well plate. The molding technique with which the gels are made within each well exposes the glass through the contact point located at the bottom of each microwell, thus facilitating adhesion of the cells to the glass surface, thereby immobilizing the spheroids during manipulations of the liquid inside the well. The coordinates of the spheroids relative to each other is identical in every well, which allows standardization and automation of the analysis obtained from the fluorescent plate imager. Due to the glass bottom and the transparency of the hydrogel, the plates are also amenable to conventional microscopy. The current design of the mold contains 19 microwells per well and therefore provides multiple replicates per condition, thereby reducing the variability and increasing the statistical power of the approach. The global oscillatory signal implies that the cells within a single spheroid respond to stimuli in a coordinated and synchronized manner^18^, a phenomenon also observed in Islets of Langerhans *in vivo*^19^. Importantly, such oscillatory activity was not observed in parallel experiments using MIN6 2D monolayers (Fig. 3 & Figure 3 – Figure supplement 1), highlighting the importance of using 3D spheroids.

The results that we obtained using MIN6 β-cell spheroids align both qualitatively and quantitatively very well with published data obtained using isolated Islets of Langerhans. For instance, we obtained EC_50_ values of 7.4 ± 0.4 mM and 7.9 ± 0.4 mM for glucose-induced calcium oscillations, consistent with previous research in which electrophysiological experiments performed on mouse Islets of Langerhans provided an EC_50_ value of 8.7 mM for glucose-induced membrane potential oscillations^20^, and EC_50_ values between 8 and 10 mM for calcium responses in human islets. Likewise, all evaluated pharmacological compounds targeting K_ATP_ channels modulated Ca^2+^ oscillations with concentration dependence profiles that match their known affinity to the target, with EC_50_ values in the nanomolar range for glimepiride and nateglinide, and low micromolar IC_50_/EC_50_ values for diazoxide and tolbutamide^21–23^. Finally, it is well established that insulin secretion from islets is driven by oscillations in β-cell cytoplasmic Ca^2+ 24^, and we indeed found that the level of secreted insulin from MIN6 β-cell spheroids correlated well with the frequency of calcium oscillations.

Finally, we used the MIN6 β-cell spheroid assay to evaluate the effect of the neurosteroid PS on glucose-induced Ca^2+^ signals and insulin release. Earlier work had shown that PS evokes a rise in intracellular Ca^2+^ and enhances glucose-induced insulin release from mouse pancreatic islets and in INS-1 human insulinoma cells, and this effect was largely attributed to PS-induced activation of TRPM3 ^25, 26^. We found that PS robustly increases the frequency of glucose-induced Ca^2+^ oscillations in MIN6 β-cell spheroids, associated with a significant potentiation of insulin release. The effects of PS on Ca^2+^ signals and insulin release were only partly abrogated in the presence of the TRPM3 antagonist isosakuranetin, suggesting that TRPM3 may not be the only molecular target for PS in these spheroids. In these experiments, PS and ISN were applied simultaneously, meaning that a kinetic offset between TRPM3 activation by PS and the establishment of effective channel blockade by isosakuranetin may have contributed to the incomplete inhibition of Ca2+ oscillations and insulin release observed in the presence of the antagonist.

While the present study focused on six pharmacological compounds spanning two mechanistically distinct ion channel targets, namely K-ATP channel modulators and TRPM3 modulators, the platform is designed to accommodate a substantially broader compound panel. Extending the assay to include modulators of additional ion channel classes relevant to β-cell physiology, such as voltage-gated Ca2+ channel blockers like verapamil or nimodipine, represents a natural next step in demonstrating the full versatility of the platform for high-throughput drug discovery applications. Furthermore, validation of the platform using alternative insulinoma cell lines, such as INS-1 cells, would provide additional evidence of its generalizability across different β-cell models and further strengthen its utility as a broadly applicable screening tool.

In conclusion, we presented a straightforward method for the formation and functional characterisation of up to 1824 β-cell spheroids on a single 96-well plate, and illustrate how the method can be used as a platform to identify small-molecules that modulate insulin secretion. The platform is easily amenable for further modifications, for instance by co-culturing with other cell types found in the Islets of Langerhans (α-cells, δ-cells and PP cells) to enhance the physiological relevance of the assay, or by the use of patient-specific stem cell-derived organoids for personalized medicine. The uniform shape and positioning of individual 3D spheroids in the multi-well plate format may also find applications in other fields of life sciences where the use of 3D structures for screening purposes is of importance.

A current limitation of the platform is its optimization for the MIN6 insulinoma cell line, which, despite its widespread use in β-cell research, differs from primary pancreatic beta cells in several important respects, including glucose sensitivity and secretory capacity. Initial attempts to adapt the protocol to hiPS-derived beta cells were unsuccessful, likely due to insufficient cellular viability following transfer to the micropatterned plate format. Optimizing the platform for use with primary beta cells or hiPS-derived beta cells therefore represents a priority for future development and would substantially enhance the physiological relevance of the assay. An additional avenue for optimization is the introduction of perfusion-induced flow, which has been shown to improve viability, insulin secretory function, and morphological integrity of 3D β-cell tissues under dynamic culture conditions ^27, 28^. Finally, while the current platform operates in a 96-well plate format providing up to 1824 individual spheroid measurements per plate, adaptation to higher density formats such as 384-well or 1536-well plates would represent a critical step towards full compatibility with industrial high-throughput screening pipelines, and is therefore identified as an important direction for future development of the platform.

## Experimental procedures

### Mold design and characterization

Micromachined 316L stainless steel molds were obtained from high-resolution CNC machining according to our computer-aided designs (CADs). The designs, consisting of molds slightly smaller than the footprint of a 96-well plate, contain 19 micro-pillar structures.

### Hydrogel substrate preparation

The wells of the glass bottom 96-well plate (P96-1.5H-N, Cellvis Inc.) were silanized to ensure adequate bonding of the PEG to the glass. 20 µL of gel solution was placed within the well and 10 μL was added on top of the mold, which was then placed in the well. The gel solution was briefly exposed to 365 nm UV light to induce polymerisation. The mold was unmolded, revealing a micropatterned hydrogel, and the well was immediately rinsed with distilled water at least three times before being submerged in water. Prior to cell seeding, the plate was sterilized using UV radiation for at least 20 minutes and sealed with a sterilized PCR film (ThermoFisher, USA). The protocol and associated hydrogel plates were developed in-house at 4Dcell (SmartSphero Plates).

### Cell culture

Mouse insulinoma 6 (MIN6) cells were grown in high glucose (25 mM) Dulbecco’s modified Eagle’s medium (DMEM) (11965092, ThermoFisher, USA) supplemented with 10% fetal bovine serum (FBS), 2% penicillin-streptomycin, 2% Glutamax, 1% sodium pyruvate and 0.001% 2-Mercaptoethanol. MIN6 cells seeded into the 96-well plates were harvested from a T-25 flask at passages 10-20 with an 80% confluency.

### β-cell spheroid formation

The PCR-film was removed, and the storage solution was aspirated from each well. The wells were then washed with PBS twice and incubated at 37°C and 5% CO_2_ with the gels submerged in 200 µL of cell culture medium per well. MIN6 cells were dissociated with TrypLE Express Enzyme (ThermoFisher, USA). After cell counting, a cell suspension was prepared in cell culture medium with a cell density of 570.000 cells per mL. An aliquot of 100 µL of cell suspension was added to each well of the 96-well plate to achieve a density of 3000 cells per microwell. The plate was then incubated at 37°C and 5% CO_2_ for 30 minutes for the cells to sediment into the microwells. An additional 150 µL of medium was added to each well before leaving the plate incubated at 37°C and 5% CO_2_ overnight. The next day, the spheroids were evaluated for correct morphology and size uniformity using brightfield optical microscopy.

### Compound plate preparation

Test compounds were prepared to the correct concentration and aliquoted into a clear 96-well plate with conical bottom (651101, Greiner Bio-One International GmbH, Austria) in accordance with the protocol for the µCELL FDSS fluorescence plate imager. All test compounds were of analytical standard or certified reference material purity (>96%) and obtained from Sigma-Aldrich (St. Louis, Mo). Stock solutions were prepared in dimethyl sulfoxide (DMSO) and stored at −20°C. Appropriate volumes of the stock solutions were added to the same KREBS solution used for the baseline measurement (NaCl 150 mM, KCl 6 mM, CaCl_2_ 2 mM, MgCl_2_ 1.5 mM, HEPES 10 mM, Glucose 3 mM, pH 7.4) to achieve the desired final concentrations in the compound plate.

### Ca^2+^ imaging

Medium was removed from the wells 18 to 24 hours after seeding and 200 µL of a 10μM Cal-520 AM (AAT Bioquest, USA) solution with 0.04% Pluronic F-127 (AAT Bioquest, USA) was added to each well and the plate was incubated at 37°C and 5% CO_2_ for 1 hour. The wells were then washed with KREBS solution (NaCl 150 mM, KCl 6 mM, CaCl_2_ 2 mM, MgCl_2_ 1.5 mM, HEPES 10 mM, Glucose 3 mM, pH 7.4) twice and incubated in 200 µL of the same KREBS solution for 30 minutes at room temperature to allow esterases to cleave the lipophilic blocking groups. The KREBS solution was removed and a final volume of 120 µL 3 mM glucose KREBS solution was placed in each well. The plate was then placed in a functional drug screening system (µCELL FDSS, Hamamatsu Photonics) fluorescence plate imager with integrated dispensing head. The automated instrument dispensed 15 µL per well of a compound plate into the assay plate according to a predetermined protocol and detects fluorescence across the plate simultaneously at a resolution sufficient to yield 4 pixels per spheroid. Ca^2+^ flux signal was captured continuously, at a sampling rate of 1 Hz, by a cooled CCD camera during the entire protocol. The excitation wavelength was set to 470 nm and the emitted light passed through a 530 nm bandpass filter. Prior to addition of the solutions from the first compound plate, a baseline was recorded for a minimum of 5 minutes. All measurements were performed at 37°C. The acquired image sequences of the fluorescence signal were exported in TIFF format for further analysis.

### Insulin detection

After 1 hour of exposure to increased levels of extracellular glucose and/or compounds, between 15 µL to 100 µL of supernatant was aspirated from the assay plate and deposited into a separate 96-well plate using the dispensing head of the µCELL FDSS. This plate was sealed with PCR film and stored at −20°C until analysis. Insulin levels in the supernatant were quantified using the Mercodia Ultrasensitive Mouse Insulin ELISA kit (Mercodia AB, Sweden) in accordance with the manufacturer’s protocol. Fluorescence values of the plate were recorded using a Flexstation 3 Microplate Reader (Molecular Devices, USA) and insulin concentrations were calculated via cubic spline regression based on the fluorescence of the calibrator solutions.

### Data analysis

Regions of interest (ROIs) were selected to contain the fluorescence signal of one spheroid per ROI using Fiji ImageJ 1.52i. The resulting signal for each individual spheroid was then analysed using a Python script (v3.10.11). The background signal was defined as the average of the pixels directly bordering the ROI and was subtracted from the ROI signal. The resulting signal was then smoothed applying a Savitzky-Golay filter with a polynomial order of 10 and a window length of 60^29^. Next, a multi-polynomial fit was applied to approximate the baseline of the signal using the IModPoly method from the BaseLineRemoval Python-module, utilizing a polynomial degree of 10^30^. The baseline was subtracted from the original fluorescence signal and peaks were identified with the find_peaks function from the SciPy.Signal data processing Python library^31^. Minimum peak height was set to a multiple of the standard deviation of the signal during basal levels of extracellular glucose (specified in-text), minimum width was set at 10 seconds and a minimal horizontal distance between peaks centres of 60 seconds. The normalized area under the peaks (AuP) was defined as the sum of the fluorescent signal for each timepoint after baseline removal within the given time window, normalized over the amplitude of the control stimulus (50mM KCl).

### Statistical analysis

Analysis was performed using Origin 9.0 (OriginLab Corporation) and GraphPad Prism 10.0 (GraphPad Software). All numerical results are expressed as mean ± standard error of the mean (SEM). EC_50_ and IC_50_ values were determined using a nonlinear regression based on the Hill equation. Assay quality was assessed by calculating the Z-factor using the Ca^2+^ oscillation frequency at 3 mM glucose as the negative control and at 20 mM glucose supplemented with 50 µM pregnenolone sulphate as the positive control.

## Data availability

The data that support the findings of this study are available from the corresponding author upon reasonable request.

## Acknowledgements

We kindly thank M. Benoit and A. Janssens for their technical support and expertise at KU Leuven. This project has received funding from the European Union’s Horizon 2020 research and innovation programme under the Marie Sklodowska-Curie grant agreement No. 955643. This work was further supported by grants from the Research Foundation — Flanders (FWO; G0B9520N and G0B7620N to T.V.), the Queen Elisabeth Medical Foundation for Neurosciences (to T.V.) and the VIB to (T.V.). Research infrastructure was funded by the KU Leuven Research Council (AKUL/19/34).

