## Supplementary figures and images for "A high-throughput assay for the measurement of Ca^2+^-oscillations and insulin release from uniformly sized β-cell spheroids"

### Supplemental information

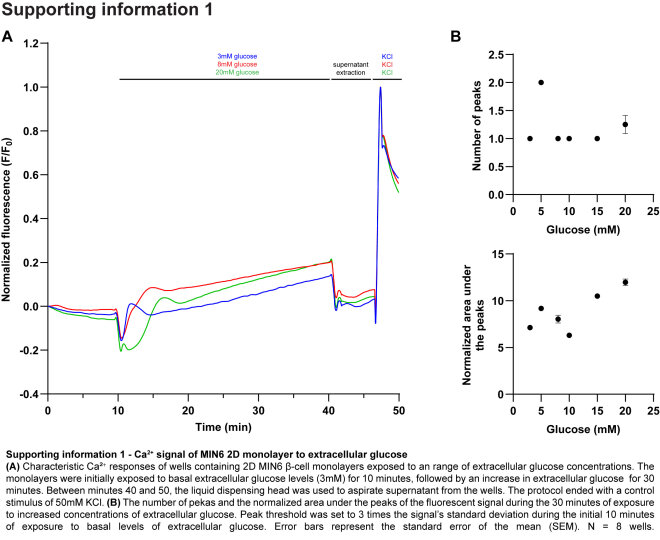
